# Ribosome clustering and surface layer reorganization in the microsporidian host-invasion apparatus

**DOI:** 10.1101/2023.05.31.543061

**Authors:** Himanshu Sharma, Nathan Jespersen, Kai Ehrenbolger, Lars-Anders Carlson, Jonas Barandun

**Affiliations:** Department of Molecular Biology, The Laboratory for Molecular Infection Medicine Sweden (MIMS), Umeå Centre for Microbial Research (UCMR), Science for Life Laboratory, Umeå University, 90187 Umeå, Sweden; Department of Medical Biochemistry and Biophysics, The Laboratory for Molecular Infection Medicine Sweden (MIMS), Wallenberg Centre for Molecular Medicine, Umeå Centre for Microbial Research (UCMR), Umeå University, 90187 Umeå, Sweden

**Keywords:** Microsporidia, Polar tube, Cryo-electron tomography, Ribosome, Polar tube proteins

## Abstract

During host cell invasion, microsporidian spores translocate their entire cytoplasmic content through a thin, hollow superstructure known as the polar tube. To achieve this, the polar tube transitions from a compact spring-like state inside the environmental spore to a long needle-like tube capable of long-range sporoplasm delivery. The unique mechanical properties of the building blocks of the polar tube allow for an explosive transition from compact to extended state and support the rapid cargo translocation process. The molecular and structural factors enabling this ultrafast process and the structural changes during cargo delivery are unknown. Here, we employ light microscopy and in situ cryo-electron tomography to visualize multiple ultrastructural states of the polar tube, allowing us to evaluate the kinetics of its germination and characterize the underlying morphological transitions. We describe a cargo-filled state with a unique ordered arrangement of microsporidian ribosomes, which cluster along the thin tube wall, and an empty post-translocation state with a reduced diameter but a thicker wall. Together with a proteomic analysis of endogenously affinity-purified polar tubes, our work provides comprehensive data on the infection apparatus of microsporidia and demonstrates that ribosomes are efficiently transported through polar tubes in a spiral-like parallel arrangement.

## Introduction

Microsporidia are a group of obligate intracellular parasites that infect hosts across the animal kingdom (Troemel *et al*, 2008; Fries *et al*, 2006; Palenzuela *et al*, 2014; Yachnis *et al*, 1996). To propagate from host to host, these fungi-like pathogens form stable environmental spores of prokaryotic size (Han & Weiss, 2017). In addition to their small size, these parasites exemplify reductive evolution in eukaryotes (Nakjang *et al*, 2013; Jespersen *et al*, 2022b). The deletion of many genes considered essential for eukaryotic function has produced some of the smallest known genomes of the kingdom (Corradi, 2015), streamlined biochemical pathways (Wadi & Reinke, 2020), and minimized cellular macromolecular complexes. Recent structural insights into microsporidian ribosomes and proteasomes bound to dormancy factors in the extracellular spore stage have highlighted the importance of mechanisms to enter and exit dormancy (Ehrenbolger *et al*, 2020; Jespersen *et al*, 2022a; Barandun *et al*, 2019; Nicholson *et al*, 2022). This trend of reductive evolution might have emerged from an obligate intracellular lifestyle, while the atypical regulatory factors support pathogen survival in nutrient-limiting environments (Heinz *et al*, 2012).

While a general trend for reductive evolution is pervasive, microsporidia have also evolved specialized mechanisms for invading and hijacking host cell systems (Xu & Weiss, 2005; Han *et al*, 2019). These mechanisms include, for example, an expanded repertoire of nucleotide transporters to steal energy and metabolic precursors from host cells (Dean *et al*, 2018; Major *et al*, 2019). However, the most drastic invention and specialization is the polar tube, a microsporidia-specific organelle used for host invasion. The polar tube comprises at least six polar tube proteins (PTPs) that may localize to its outermost layer or the terminal tip (Han *et al*, 2017; Weidner, 1976; Lv *et al*, 2020), but the structure and organization of PTPs and the exact composition of the tube is still a mystery. Inside the spore, this organelle adopts a spring-like arrangement and is referred to as the polar filament (Frixione *et al*, 1992; Han *et al*, 2020; Jaroenlak *et al*, 2020). Polar filament coils can range from a couple to dozens, depending on the organism (Xu & Weiss, 2005). Upon exposure to the right environmental stimuli, the spores germinate. This includes an explosive firing at the apical pole of the spore that transforms the polar filament into the extended, tube-like state referred to as the polar tube.

During germination, the entire infectious cellular content, known as the sporoplasm, is pushed through the narrow polar tube and delivered into or close to the host cell (Franzen, 2004). These events, including tube eversion and passage of sporoplasm cargo through this constricted tube, are extremely fast and occur within 2 seconds (Jaroenlak *et al*, 2020). To achieve successful firing, sporoplasm delivery, and host infection, the polar tube and the cargo undergo drastic remodeling during this discharge (Jaroenlak *et al*, 2020; Troemel & Becnel, 2015). These swift ultrastructure changes in the polar tube evince its extraordinary mechanical properties likely conferred by its components, the PTPs. However, the mechanisms driving polar tube firing, sporoplasm discharge, and the dynamics of polar tube eversion are not well understood (Jaroenlak *et al*, 2020; Chang *et al*, 2023). Further, the swift nature of germination events creates challenges in capturing polar tube dynamics and contributes to the generally understudied nature of these parasites. Additionally, most *in vitro* studies on the microsporidian infection apparatus using electron microscopy (EM) employ denaturing purification of polar tube components or have relied on non-native staining and resin embedding methods, thus limiting their overall resolution.

To overcome these challenges, we employed light microscopy and cryo-electron tomography (cryo-ET) to examine the dynamics and ultrastructure remodeling of the *Vairimorpha necatrix* polar tube during germination in a near-native environment. We capture snapshots of the infectious sporoplasm seen as arrays of organized ribosomes and densely packed cargo passing through the polar tube. Further, two distinct states of the extruded polar tube, discernible in the protein layer lining its outer wall, are also described. We further characterize the outer proteinaceous layer of the tube using affinity purification and mass spectrometry, thus unraveling potential new protein-protein interactions of the PTPs. Overall, our results shed new light on the events underpinning cargo delivery by the polar tube and uncover protein factors that may assist host invasion.

## Results

### Cryo-ET captures assorted states of the polar tube and cellular content during cargo delivery

The microsporidian infection apparatus rapidly transitions from a tightly packaged polar filament to an extended polar tube. The dynamics of germination and polar tube length have been studied in three human-infecting microsporidia; however, spore firing is known to vary between species, and firing efficiencies can also vary greatly even for uniform-looking spores (Jaroenlak *et al*, 2020). We set out to identify and optimize conditions for *V. necatrix* germination and quantify the dynamics of tube firing in this agriculturally important parasite of Lepidoptera (Mitchell & Cali, 1994; Kurtti *et al*, 1990). We used live microscopy to record 53 germination events, which enabled us to measure the polar tube length and firing velocity (**Supplementary Fig. 1**). We observed that polar tubes could attain a maximum length of ∼142 µm, a mean length of 113 µm, and extend with a mean maximum velocity of ∼281 µm/s. Similar to the previous observations in distant microsporidia (Jaroenlak *et al*, 2020), *V. necatrix* everts its tube to the maximum length in less than one second and expels the sporoplasm from the tip of the nascent polar tube. Additionally, tube firing follows a typically observed triphasic mode where the tube undergoes observable states of elongation, followed by stasis during sporoplasm passage and a refractory period where the tube shortens after cargo emergence (Jaroenlak *et al*, 2020) (**Supplementary Fig. 1**). Akin to tube remodeling events, the sporoplasm itself also swiftly transforms from a restricted spore state to an extremely extended conformation during extrusion into a circular shape upon emergence (**Supplementary Fig. 1a**), and these events likely impose immediate reorganization of subcellular structures.

The optimal conditions for polar tube firing were used to cryogenically preserve on-grid germinated spores (**Supplementary Fig. 2a**) for *in situ* characterization of polar tubes and cargo using cryo-ET. On-grid germination was optimized with spores consistently displaying high germination efficiencies. Spores were applied onto cryo-grids immediately after resuspension in the germination buffer, followed by blotting and plunge freezing. Various grid types were screened during freezing attempts, and polar tubes frequently interacted strongly with the regular grid support regions while avoiding thin carbon films or holes. To overcome this, we utilized lacey carbon grids with an ultra-thin carbon film where the variability in hole size and support mesh increased the chance of trapping sections of the tubes on larger thin-carbon areas. Subsequently, we collected 50 tilt series from different sections of polar tubes (**Supplementary Table 1**), of which 45 were suitable for tomogram reconstruction using the IMOD package (Kremer *et al*, 1996). The reconstructed tomograms were denoised using IsoNet (Liu *et al*, 2022) and auto-segmented using CNN in Eman2 (Chen *et al*, 2017) to better visualize the spatial organization of the cargo and polar tubes. This allowed us to capture a wide range of polar tube ultrastructural states, with differences in diameter, outer layer thickness, and interior composition, most likely resulting from the differential passage of reorganized sporoplasm.

The diameter of the analyzed polar tubes ranges from 60 nm to 190 nm depending on their internal content, suggesting we captured a range of different phases during germination or after sporoplasm translocation (**Fig. 1**). Of note was an apparent correlation between the germination phase and the thickness of the tube wall, which is composed of a lipid bilayer (pink arrows, **Fig. 1** & **Supplementary Fig. 2**), flanked by an outer layer of polar tube proteins (light blue and magenta arrows, **Fig. 1** & **Supplementary Fig. 2**). Polar tubes filled with dense cellular cargo (*PTcargo*) (**Fig. 1a-c**) had diameters of more than 120 nm and are sheathed by a thin layer; in contrast, the electron-lucent or empty tubes (**Fig. 1d, e**) have a diameter of significantly less than 120 nm but were enveloped by a thick outer layer (*PTempty*) (**Supplementary Fig. 2b**). Most interestingly, across all our tomograms, we observed different internal cargos, including, electron-dense material inside membranous compartments that could be attributed to organelles like the nucleus, randomly oriented or highly ordered large molecular complexes such as ribosomes and proteasomes, and empty vesicles or empty tubes. The electron-dense cellular material seemed free-flowing or sometimes enveloped inside vesicle-like organelles (orange arrows, **Fig. 1a, Supplementary Fig. 2d**), while distinct densities corresponding to macromolecular complexes were randomly distributed or arranged in a very regular fashion (**Fig. 1b, c**). Vesicle-containing tubes were reminiscent of the frequently observed tube-inside-tube architecture of the polar tube (Takvorian *et al*, 2020). Notably, macromolecular complexes were completely missing from *PTempty*, which mostly housed empty-looking vesicles (dark blue arrows, **Fig 1d** & **Supplementary Fig. 2c**) or no vesicles (**Fig 1e** & **Supplementary Fig. 2c**). Collectively, these tomograms capture the myriad heterogeneous states the polar tubes and the cellular cargo adopt immediately upon firing.

**Figure 1.**
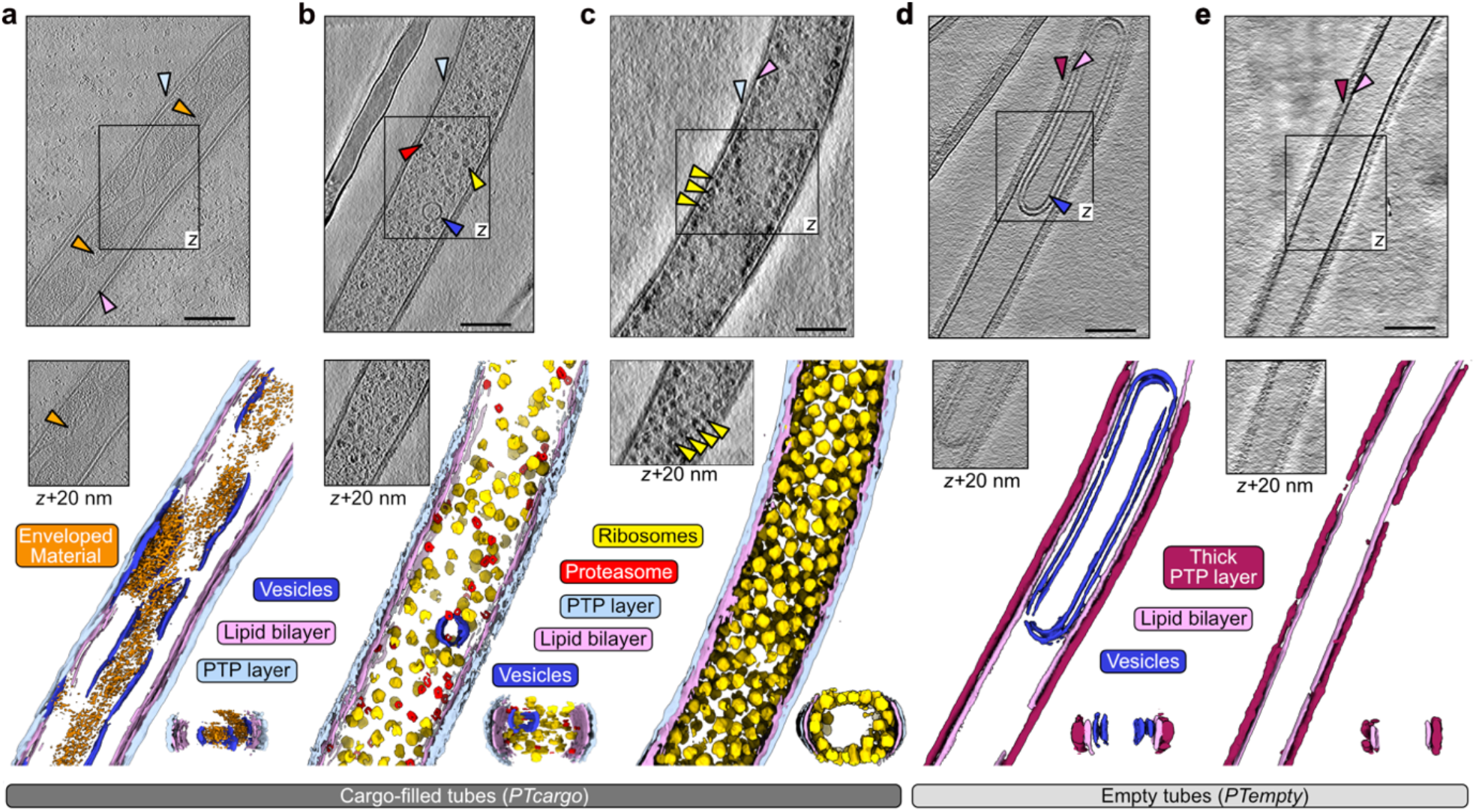
Ultrastructural states of the polar tubes during or after delivering cellular cargo. (**a-e**) Slices through cryo-tomograms of germinated polar tubes above the respective neural network aided 3D segmentation. The central slice of a tomogram is represented at the top in each panel, as seen from the z-axis view. An additional section from the same tomogram is marked by a boxed region and shown below at axis z + 20 nm. The neural network-aided 3D segmentations corresponding to each tomogram are presented in the lower panel. Selected regions of interest in the tomogram slices, such as cellular complexes (a-c), the polar tube layer (a-e), or empty vesicles (d), are indicated with arrows colored as the labels and segmentations below. The black scale bars correspond to 100 nm.

### Regular clustering of ribosomes in the cargo-filled polar tubes

In five out of the 45 cryo-ET reconstructions, we captured sporoplasm sections with an unusually high concentration of ribosome-like particles (**Fig. 2** and **Supplementary Fig. 2e-f**). The particles were arranged in an array-like pattern (**Fig. 2a**), wherein their parallel alignment formed a right-handed helix traversing the entire length of the reconstruction (**Fig. 2b**). These arrays were prominent in the distal sections of the tomograms and the y-axis view of the tube cross-sections, indicating the spirals line the inner wall of the tube (**Fig. 2a, b)**. To confirm their identity, we performed reference-free subtomogram averaging on the spirally arranged ribosome-like particles and obtained a low-resolution map with clear ribosome features (**Fig. 2c, Supplementary Fig. 3a**). This map superimposes well with the published high-resolution structure of the *V. necatrix* ribosome (Barandun *et al*, 2019) (**Fig. 2c, Supplementary Fig. 3b**). We could unambiguously identify features of the 40S and 60S ribosomal subunits thus affirming that the spirally arranged particles are indeed ribosomes (**Fig. 2c**). Among these features, density likely corresponding to parts of the well-conserved L7/L12 stalk region was also observed. To further understand the overall organization of the ribosome spirals, we measured the inter-particle distance between the four nearest neighboring macromolecules (**Fig. 2d, e**). The nearest particle pairs approach 200 å, with a mean interparticle distance of 220 å (d1), spread uniformly across the tube. In contrast, the other adjacent ribosomes (d2, d3) are approximately 300 å distant, on average. An inter-ribosome distance of 220 å matches well with the observed 100S hibernating ribosome dimer from *Spraguea lophii* (McLaren *et al*, 2022), suggesting that the parallel spiral is composed of repetitive sets of hibernating ribosomes organized into an almost crystalline-like arrangement inside the tube.

**Figure 2.**
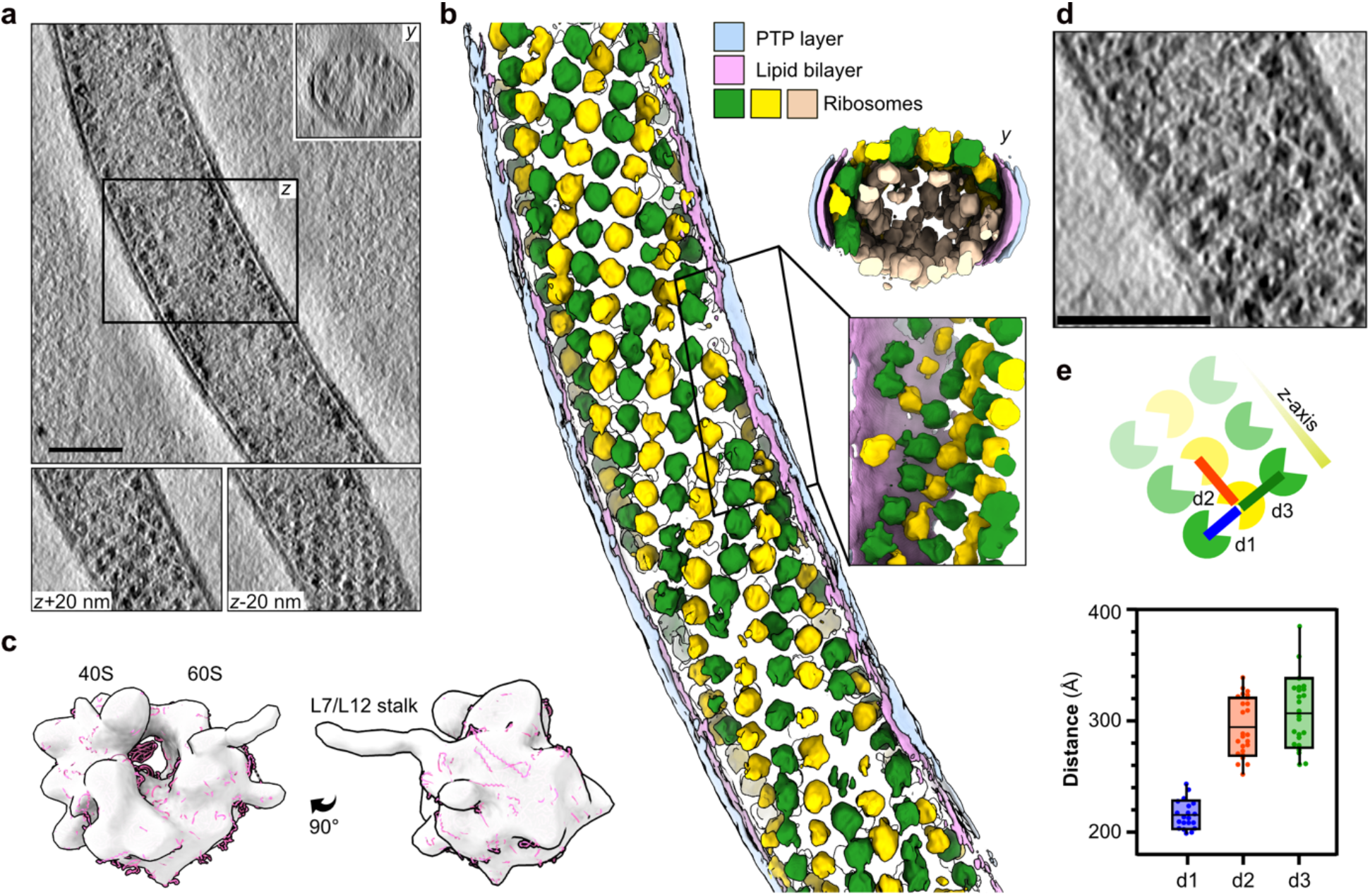
Spiral-like arrays of ribosomes clustering in proximity to the tube wall. (**a**) Slices through a cryo-tomogram and the corresponding neural network aided 3D segmentation (**b**) of a polar tube filled with clustered ribosome arrays. The segmentation presents ribosomes in yellow, green, or beige. The lipid bilayer is shown in pink, while the outermost PTP layer is shown in blue. A y-axis view of the tomogram, as well as slices at z+20 nm and z-20 nm, are shown in the inset. A similar view along the y-axis is depicted with the segmented tube. The scale bar is 100 nm. (**c**) The subtomogram average of clustered particles picked from cargo-filled polar tubes. The map is lowpass filtered to at a resolution of 50 å and is superimposed with the structure of the *V. necatrix* ribosome (PDB ID: 6RM3, magenta). (**d**) A zoomed-in view of the ribosome array packing inside the polar tube. The scale bar is 100 nm. (**e**) A schematic representation of panel (d) with the inter-ribosome distances (d1-d3) indicated (top) and plotted (bottom). The scatter plot depicts the average distance and distribution between particle centers. Each dot represents a ribosome pair chosen for measuring the distances. The distance d1 corresponds to two close particles, while d2 and d3 are the more distant.

### Remodeling of the polar tube protein layer during cargo translocation

Visualizing polar tube sections in different states allowed us to investigate the ultrastructural organization of the outer layer in empty and filled tubes (**Fig. 1a-e, and Fig. 3a**). Since the presence or absence of cargo significantly affected the tube diameter, we assumed this could also affect the structure of the outer layer. To investigate this, we compared the coats of cargo-filled and empty tubes by manually measuring the thickness of the individual layers and by subtomogram averaging particles picked on the tube wall from both states. In our tomograms, we could resolve two individual layers where the inner layer had two electron-dense leaflets suggesting it be a lipid bilayer membrane. In contrast, the outer layer was less electron-dense, and we hypothesize that this layer is composed of the PTPs, which would be consistent with previous studies (Xu *et al*, 2004). The presence of an outer protein-composed tube wall also agrees well with immunolabeling studies where fluorescently labeled antibodies against individual PTPs localized to the outer sheath of the tube wall (Han *et al*, 2017). We surveyed the thickness of the inner lipid bilayer membrane and the outer protein/PTP-composed layer (**Fig. 3b)** across several tomograms and compared the measured values between empty and cargo-filled tubes. The thickness of the lipid bilayer was, approximately 30 to 40 å, relatively consistent between both states (**Fig. 3c)**. In strong contrast, the PTP layer thickness varied significantly between the two polar tube states where the outer layer was around 50 å thick in all measured regions of *PTcargo* but ranged from 100 to 150 å in *PTempty* (**Fig. 3c)**.

**Figure 3.**
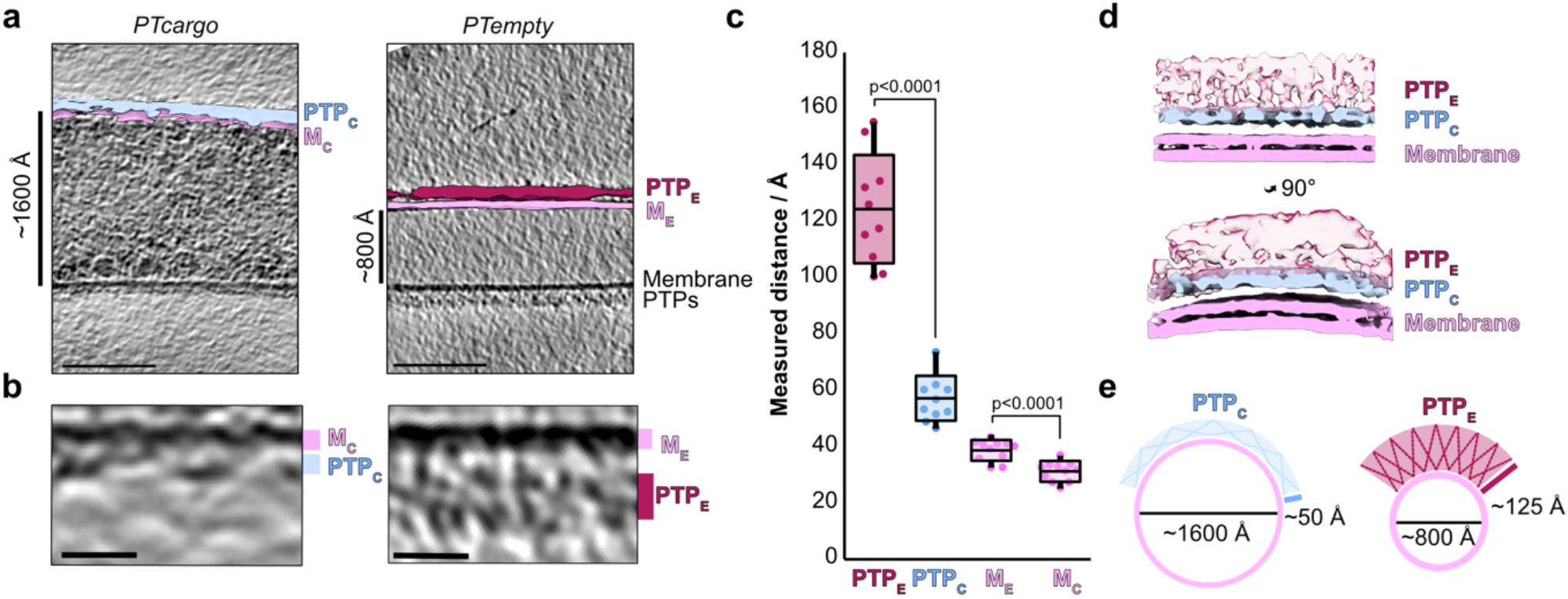
Visualizing polar tube wall features from cargo-filled and empty tubes. (**a**) Central slices through representative cryo-tomograms from cargo-filled and empty polar tubes. Segmentations of the inner and outer layers are superimposed onto one side of the tube in both tomogram slices. The scale bar is 100 nm. (**b**) Zoomed in sections of the tube wall from (a). The scale bar is 10 nm. (**c**) A plot depicting the thickness of each PTP layer and membrane bilayer, as measured across various tomograms, for cargo-filled and empty polar tubes. The p-values of two-tailed, unpaired Student’s *t*-test analyses are shown above the compared plots. (**d**) Overlay images of subtomogram averages from related views of cargo-filled and empty polar tubes (same view rotated 90° around the Y-axis). Map regions are colored as in (a), and the PTP layer from empty tubes (PTP_E_) is presented as partially transparent. (**e**) A schematic representation of polar tube and PTP layer remodeling during cargo movement.

Further, we performed subtomogram averaging with sections extracted from the outer layer of the empty and cargo-filled tubes. The averages derived from the two ultrastructural states of the tube resolved the membrane into two density layers as expected for a lipid bilayer (**Fig. 3, Supplementary Fig. 3c**), in good agreement with the known thickness of membranes (Tristram-Nagle & Nagle, 2004). In addition, the subtomogram averages reiterate the differences observed in PTP layer thickness (**Fig. 3d)**. This suggests the protein wall has unique mechanical properties that allow it to stretch out into a thin layer when cargo traverses through the tube and contract again once the sporoplasm is ejected at the tip (**Fig. 3e**). These observations collectively suggest the importance of a dynamic PTP layer that undergoes major structural remodeling to facilitate polymorphic states of the polar tube during cargo delivery (**Fig. 3e**).

### Native composition of the polar tube protein layer

Intrigued by the large-scale remodeling of the PTP layer, we focused on understanding its composition. For this, we discovered a stretch of eight proximate histidines, serendipitously located in the known polar tube component PTP3 from *V. necatrix* (**Fig. 4a)**. Generally, PTP3 is the largest among known polar tube components and is predicted to harbor a cleavable signal peptide and stretches of disordered regions. PTP3 antibodies localize to the entire length of the polar tube except for the tip (Peuvel *et al*, 2002). This suggests that PTP3 is part of the protein scaffold of the tube, which makes it an ideal candidate for endogenous compositional analysis. We affinity-purified endogenous PTP3 and its potential interaction partners for protein identification analysis using mass spectrometry (**Fig. 4b, Supplementary Table 2**). In this sample, PTP3 was significantly enriched and the most abundant protein (**Fig. 4c-e**), confirming the utility of the endogenous poly-histidine patch for affinity purification. The known polar tube members PTP1 and PTP2 were also captured and present in our pull-down, where PTP2 was the second most enriched protein among all hits (**Fig. 4e**). This suggests direct or indirect interaction of PTP1 and PTP2 with PTP3, which agrees with previous yeast two-hybrid assays for PTPs from *Encephalitozoon cuniculi* (Bouzahzah *et al*, 2010). However, neither PTP5 nor the polar tube tip localizing PTP4 (Han *et al*, 2017), could be detected, suggesting the absence of direct or stable interactions with PTP3. This is also consistent with recent observations that PTP3 is not present at the tip of the tube (Fayet *et al*, 2023).

**Figure 4.**
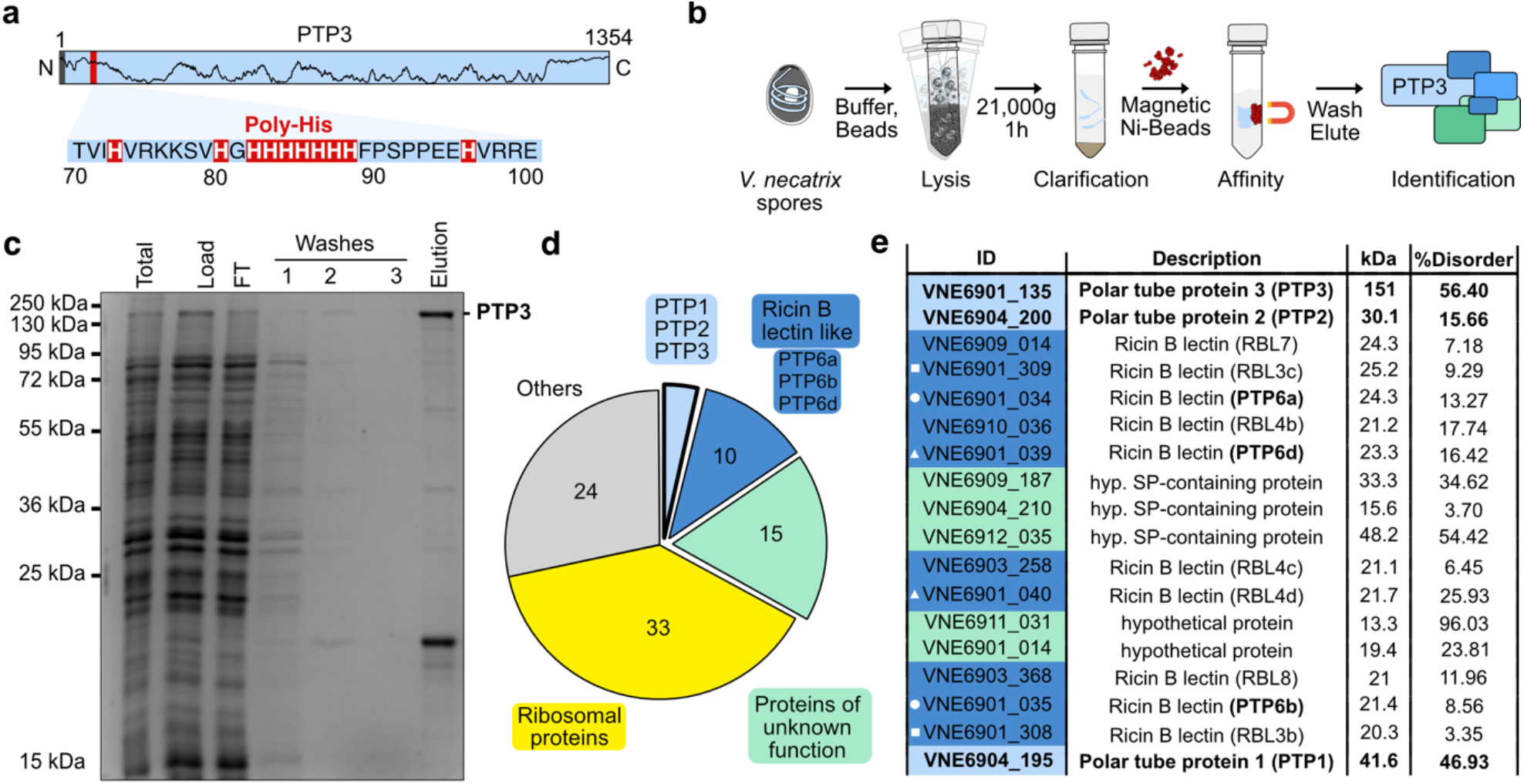
Endogenous PTP3 pulldown. (**a**) Representation of the 1354 amino acids long *V. necatrix* PTP3. Marked are the N-terminal predicted signal peptide (highlighted in grey), the histidine stretch (red area zoomed in with sequence) that was used for purification, and the disorder prediction (black line). (**b**) Schematic representation of the endogenous purification of PTP3. (**c**) The different purification steps shown in (b) have been analyzed on an SDS PAGE. (**d**) A pie chart showing the distribution of all mass spec hits detected in the elution sample, shown in (c), classified using functional and structural annotation of the *V. necatrix* genome (*manuscript in preparation*). (**e**) Mass spec hits from the classes: PTPs, RBLs and proteins of unknown function, are sorted based on the number of significantly detected unique peptides (high to low). Gene IDs are presented with colors for the respective classes, along with names for the encoded proteins, molecular weights, and structurally disordered regions predicted using PrDos (Ishida & Kinoshita, 2007). The white symbol in front of the gene ID indicates close genomic localization.

In addition to the known polar tube members, other enriched proteins in our mass spectrometry analysis suggest additional factors may constitute the microsporidian infection organ. The remaining hits were segregated based on direct functional assignment or the presence of signature structural features (**Fig. 4d**). A significant class with Ricin-B lectin domain-containing proteins (RBLs) accounts for more than 10% of all hits. After PTP3, several RBLs were the next most enriched proteins in the pulldown, and may correspond to proteins migrating around 20 kDa in the gel (**Fig. 4c)**. RBLs contain the B-chain of the Ricin glycoprotein and enable interactions with β-linked galactose or N-acetylgalactosamine containing glycans (Cummings & Etzler, 2009), and assist infection by promoting host cell adherence (Prybylski *et al*, 2022; Liu *et al*, 2016). We noted that the tandem genomic location for some of the detected RBL encoding genes agrees well with the previously observed synteny in microsporidian genomes (Campbell *et al*, 2013) and may indicate the presence of dimers (e.g. VNE6901_034/035, VNE6901_039/040, VNE6901_308/309). Interestingly, among these RBLs, VNE6901_034, VNE6901_035 and VNE6901_039 possess sequence and structure homology to *Nosema bombycis* PTP6 **(Supplementary Fig. 4)**, also known to localize throughout the polar tube (Lv *et al*, 2020), suggesting the presence of PTP6 isoforms and their interaction with PTP3.

Additionally, several proteins of unknown function (>25%), containing spore wall-like proteins, proteins with signal peptides (SP), or high propensity of disordered regions, were identified (**Fig. 4e, Supplementary Table 2**). Finally, the remaining hits could be attributed to ribosomal proteins (RPs) and other proteins involved in cellular maintenance (**Supplementary Table 2**). Although our work discovered ribosomes that cluster along the inner PT wall (**Fig. 2**), we assume that the RP hits represent contaminations, as their high abundance and the unusual biochemical properties often lead to the identification of peptides from RPs via mass-spectrometry. Collectively, this mass spectrometry data, to our knowledge, provides the first endogenous compositional analysis of an affinity-purified polar tube and suggests novel interactions between RBLs and PTPs, and also uncovers new candidates that may assist polar tube structure and function.

## Discussion

Host cell invasion is an understudied aspect of the microsporidian lifecycle that initiates with polar tube firing to inoculate host cells with the infective sporoplasm (Franzen, 2004). The mechanisms enabling tube remodeling, ensuing sporoplasm translocation through an extremely constricted passage, and the state of sporoplasm during delivery remain unaddressed. Here, we probe cargo delivery by cryogenically preserving snapshots of spore germination. Congruence in polar tube firing kinetics seen in our work and previous studies (Frixione *et al*, 1992; Jaroenlak *et al*, 2020), suggest a conserved theme of sporoplasm movement upon germination (**Supplementary Fig. 1**). Our tomograms show the heterogeneous states of free-flowing macromolecular complexes, vesicles of varying electron density (**Fig. 1**) and spirally packed hibernating ribosomes (**Fig. 2**), traversing the tubes. The presence of membranous vesicles and similarly encased nucleus-like, electron-dense material suggests the maintenance of compartmentalization even during explosive events that mandate rapid cellular deformations to facilitate cargo movement in the tube. Another type of non-membranous compartmentalization is seen as long strands of highly organized ribosome spirals in fired tubes (**Fig. 2**). Similar crystalline arrays of eukaryotic ribosomes, formed as part of a stress response (Milligan & Unwin, 1982; Byers, 1971; Morimoto *et al*, 1972a) or upon chemical treatments (Kusamrarn *et al*, 1975), display a reversible-metabolically inactive state and resistance to nucleolytic degradation (Morimoto *et al*, 1972b). Collectively, akin to hibernating ribosome dimers in bacteria (Franken *et al*, 2017; Matzov *et al*, 2019), the spiral of ribosome dimers in microsporidia may aid in ribosome hibernation, further adding to the atypical translation regulation mechanisms seen in microsporidian ribosomes (Barandun *et al*, 2019; Ehrenbolger *et al*, 2020). Alternatively, these spirals could represent a transient state of polyribosomes seen in pre-spore stages of intracellularly growing microsporidia (Bacela-Spychalska *et al*, 2018) and precursors to the free hibernating ribosome dimers (McLaren *et al*, 2022). This would also provide an efficient strategy for ribosome packing, transportation, and redeployment, wherein rapidly translocating arrayed ribosomes may induce ribosome crowding (Vande Broek *et al*, 1999; Karpova & Gillet, 2018; Maheshwari *et al*, 2023) for a translation jumpstart to maximize protein synthesis rates. Similarly, compartmentalization would facilitate the immediate restructuring of sporoplasms after host cell invasion to expedite the hijacking of host functions.

Our work establishes that the polar tube undergoes large-scale structural reorganization during germination. Most dramatically, the tubes seem to nearly double in internal diameter and quadrupled tube volume in the *PTcargo* state (**Fig. 3**). Collectively, these observations suggest that both tube ultrastructure and sporoplasm reorganizations (Jaroenlak *et al*, 2020) are contemporaneous events facilitating sporoplasm delivery. Here the remodeling of a highly dynamic tube protein layer around the fluid lipid bilayer drives these large-scale ultrastructure changes (**Fig. 3**), likely in response to the hydrodynamic pressure exerted by the traversing sporoplasm. Such instances of mechanical recoil are widely reported in elastins and other membrane-localizing disordered proteins involved in mechanical and trafficking functions in cells. Interestingly, PTP1 and PTP3 and other factors identified in our native pulldown experiments (**Fig. 4**), contain a significant fraction of predicted disorder. The high structural flexibility of disordered proteins provides a large conformational sample space for structure and membrane curvature remodeling under tension (Rauscher & Pomès, 2017; Yuan *et al*, 2021; Fakhree *et al*, 2019). Further, the observed post-translational O-glycosylation and mannosylation of PTPs (Xu *et al*, 2004; Bouzahzah *et al*, 2010; Keohane *et al*, 1998) may further expand their protein-protein interaction repertoire, structural flexibility (Prates *et al*, 2018) and host cell attachment abilities (Hounsell *et al*, 1996; Casanova *et al*, 1992; Lin *et al*, 2020). Additionally, our native purifications suggest an association of PTPs with several carbohydrate-binding RBLs that promote host-cell attachment in microsporidia and other intracellular pathogens (Prybylski *et al*, 2022; Sardinha-Silva *et al*, 2019; Nogueira *et al*, 2016; Petri Jr *et al*, 2002). Further, the presence of signal peptides-containing hypothetical proteins suggests extracellular localization and potential PTP interaction to form the outer, flexible layer of the polar tube.

In summary, our study provides structural insights into the germinated microsporidian polar tube, revealing a spiral array-like arrangement of ribosomes, two distinct structural states of the polar tube protein layer, and new potential components of this layer. Further validation and biochemical characterization of these potential PTPs and a higher resolution structure of the polar tube components will help us fully understand the mechanistic basis of tube reorganization and the role of individual PTPs and RBLs in host cell invasion.

## Methods

### Spore isolation

*V. necatrix* was cultivated and reproduced by feeding approximately 100,000 spores to fourth and fifth instar *Helicoverpa zea* larvae grown on a defined diet (Benzon Research). After three weeks at 21–25 °C, the spores were harvested. First, larvae were homogenized in water, followed by filtration through two layers of cheesecloth and subsequent filtration through a 50 μm nylon mesh. The filtrate was layered on top of a 50% Percoll cushion in a 2-ml microcentrifuge tube, and spores were pelleted by centrifugation at 1,000g for 10 minutes. The pure spores were stored at -80 °C until further use.

### Germination of *V. necatrix* spores

Purified *V. necatrix* spores were germinated via alkaline priming (Kurtti *et al*, 1990). Briefly, spores were incubated in 100 μl of 0.01 M KOH for 15 minutes at room temperature, followed by pelleting via centrifugation at 10,000g for 2 minutes. Primed spores were then germinated by resuspending in 100 μl germination buffer (0.17 M KCl, 1 mM Tris-HCl (pH 8.0), 10 mM EDTA). Germination was confirmed by light microscopy. Approximately 80% of spores germinated with the immediate addition of the germination buffer.

### Light microscopy

To examine the germination process via light microscopy, 0.1 mg of alkaline-primed *V. necatrix* spores were resuspended in 50 μl of the germination buffer. Next, 2.5 μl of spore suspension was immediately transferred to a glass slide and sealed with a cover slip. Germination occurred up to 5 minutes after resuspension in the germination buffer. Videos of polar tube firing were captured using a Nikon 90i Eclipse microscope equipped with a 40x PH2 phase-contrast objective lens and a Hamamatsu C4742-80 ORCA-ER digital camera, collecting images at 15 frames per second.

Kymographs were produced using the straightening function within the FIJI image analysis software (Schindelin *et al*, 2012). To quantify the lengths and maximal velocities, polar tube length was measured on a frame-by-frame basis in the FIJI software using the segmented line function, starting with the polar tube exit site on the germinating spore. Maximal velocity was calculated as the point of greatest change in length per frame.

### On-grid germination of spores and data collection

Alkaline-primed spores were resuspended in the germination buffer and immediately blotted onto glow-discharged Lacey Carbon 200-UT grids. Grids were blotted with a blot time of 3.5 to 4.5 seconds with a blot force of -1 using an FEI Vitrobot Mark IV (Thermo Fisher Scientific), followed by plunge freezing in liquid ethane. The vitrobot was set to 4 °C and 100% humidity throughout the blotting process. Tilt series were collected on a Titan Krios (Thermo Fisher Scientific) operated at 300 kV using a Gatan K2 BioQuantum direct electron detector at the Umeå Core Facility for Electron Microscopy. Tilt series were collected on the Tomo5 package (Thermo Fisher Scientific) using an object pixel size of 2.173 å. A variable tilt angle range of -60° to +60° with a step range of 2° or 3° was used for a dose-symmetric data collection scheme (Hagen *et al*, 2017). A nominal defocus range of -1.5 to 5 μm was utilized across the data collections, the total electron dosage was fixed at 110 - 120 e-/å^2^, and a total of 50 tilt-series were collected, of which 45 were used for analysis. Positions for tilt-series collection were chosen by visual inspection of tube regions based on a heterogenous mix of thickness and electron density of visible features. Spores and polar tube tips with ejected cargo were too electron dense for collecting tilt series and hence were excluded from data collection.

### Tilt series processing, tomogram generation and segmentation

Tilt series motion correction was done using MotionCor2 (Zheng *et al*, 2017). Tilt-series alignment was done using patch-tracking in IMOD followed by CTF correction using CTFPlotter (Kremer *et al*, 1996). Dose filtering and reconstruction were performed using the IMOD package. Tomograms were reconstructed using the weighted back projection and subsequently binned four times for analysis. Tomograms were denoised via Isonet filtering (Liu *et al*, 2022) and imported into EMAN2 for CNN-based picking and segmentation (Chen *et al*, 2017).

### Subtomogram averaging

Subtomogram averaging was carried out as schematically indicated in (**Supplementary Fig. 3**). Around 150 particles were manually picked and extracted from 4-times-binned tomograms using Dynamo (Castaño-Díez *et al*, 2012). The resulting particles were aligned and centered manually before generating a first average representing a ribosome blob. Subsequently, the inbuilt template-matching function in Dynamo was used to pick particles using a low-pass filtered initial model. Post template matching, coordinates were manually inspected and cleaned to retain picks arising from the spiral arrangement, and approximately 3000 particles were extracted for the final alignment. A spherical mask centered on the initial model was created for alignments, and alignment was performed with limiting shifts and allowing for full-azimuthal and full-in-plane rotations. Azimuthal angles of the particles in the crop table were then randomized to decrease the impact of the missing wedge, and by this process, another average was generated. Full-azimuthal rotations and limited (±60°) in-plane rotation were performed with C1 symmetry for the final averaging round. The resolutions were estimated to 49 å using the Gold-standard Fourier shell correlation with a threshold of 0.143. A similar methodology was utilized for subtomogram averaging of ribosome dimers picked along the inter-ribosome distance d1. However, averaging attempts for ribosome dimer repeatedly generated anisotropic subvolumes, likely due to a specific orientation problem.

For subtomogram averages of sections of polar tube wall, Dynamo’s inherent surface models were generated manually for each tomogram and utilized for defining initial orientation before particle extraction. Extracted particles from one tomogram of each *PTcargo* and *PTempty*, respectively, were averaged without alignment to create initial models. Subsequently, particles were extracted from all tomograms. A first round of alignment run was performed using the initial model as a template with angular sampling similar to the ribosome alignment described above, along with removing oversampled particles based on the “separation in tomogram” parameter. Particles averaged after the removal of duplicates were extracted to create a new table and used for the final alignment. Resolution estimates using the Gold-standard Fourier shell correlation with a threshold of 0.143 for *PTempty* segments stood at 26 å but could not be determined reliably for *PTcargo* segments.

### Affinity purification of PTP3 and associated proteins

To enrich a polar tube sample, 50 mg of *V. necatrix* spores were resuspended in 500 μl lysis buffer (50 mM Tris pH 8.0 at 4 °C, 150 mM NaCl, 40 mM Imidazole, 5 mM DTT, protease inhibitor cocktail consisting of PMSF, E64, Pepstatin), and DNAseI. Spores were lysed via bead beating in tubes containing Lysing Matrix E (MP Bio) in 3 × 1-min intervals in a Fast-Prep 24 (MP Bio) grinder at 5.5 m/s with 5 min break on ice in-between. The lysate was transferred into new tubes and centrifuged for 1 h at 21,000g and 4 °C. The clarified lysate was then incubated with 100 µl His Mag Sepharose Ni beads (Cytivia) rotating at 4 °C to capture and enrich *V. necatrix* PTP3 via the poly-histidine patch on its N-terminal end (**Fig 4a**). The beads were washed three times with 1 ml wash Buffer (50 mM Tris pH 8.0 at 4 °C, 150 mM NaCl, 80 mM Imidazole, 5 mM DTT) and eluted in 80 µl elution buffer (50 mM Tris pH 8.0 at 4 °C, 150 mM NaCl, 250 mM Imidazole, 5 mM DTT). The sample was analyzed on SDS-PAGE and sent for in-solution mass spectrometry analysis.

### Proteomics sample preparation

The sample was reduced with DL-dithiothreitol (DTT, 100 mM) at 60 °C for 30 min and digested with trypsin using a modified filter-aided sample preparation (FASP) method (Wiśniewski *et al*, 2009). In short, the reduced sample was transferred to a Microcon-30 kDa centrifugal filter (Merck), then washed repeatedly with 8 M Urea, 50 mM triethylammonium bicarbonate (TEAB) and once with digestion buffer (0.5% sodium deoxycholate (SDC), 50 mM TEAB). The reduced cysteine side chains were alkylated with 10 mM methyl methanethiosulfonate (MMTS) in the digestion buffer for 30 min at room temperature. The sample was repeatedly washed with a digestion buffer and digested with trypsin (0.2 µg, Pierce MS grade Trypsin, Thermo Fisher Scientific) at 37 °C overnight. An additional portion of trypsin (0.2 µg) was added and incubated for another 3 hours the next day. The peptides were collected by centrifugation, and SDC was removed by acidification with 10% trifluoroacetic acid. The sample was purified using High Protein and Peptide Recovery Detergent Removal Spin Column (Thermo Fisher Scientific) and Pierce peptide desalting spin columns (Thermo Fisher Scientific) according to the manufacturer’s instructions. The purified peptide sample was dried on a vacuum centrifuge and reconstituted in 3% acetonitrile, and 0.2% formic acid for the LC-MS/MS analysis.

### NanoLC-MS analysis and database matching

Analysis was performed on an Orbitrap Exploris™ 480 mass spectrometer interfaced with an Easy-nLC1200 nanoflow liquid chromatography system (Thermo Fisher Scientific). Peptides were trapped on an Acclaim Pepmap 100 C18 trap column (100 μm x 2 cm, particle size 5 μm, Thermo Fisher Scientific) and separated on an in-house packed analytical column (75 μm x 35 cm, particle size 3 μm, Reprosil-Pur C18, Dr. Maisch) from 5% to 45% B over 78 min, followed by an increase to 100% B at a flow of 300 nl/min. Solvent A was 0.2% formic acid, and solvent B was 80% acetonitrile, 0.2% formic acid. MS scans were performed at 120,000 resolution, m/z range 380-1500. MS/MS analysis was performed in a data-dependent manner, with a cycle time of 2 s for the most intense doubly or multiply charged precursor ions. Precursor ions were isolated in the quadrupole with a 0.7 m/z isolation window, with dynamic exclusion set to 10 ppm and a duration of 30 seconds. Isolated precursor ions were fragmented with higher energy collisional dissociation (HCD) set to 30%, AGC target was set to 200%, and the maximum injection time to 54 ms. Data analysis was performed using Proteome Discoverer version 2.4 (Thermo Fisher Scientific). The raw data was matched against *V. necatrix* proteome derived from a high-quality genome assembly *(manuscript in preparation)* using Mascot 2.5.1 (Matrix Science) as a database search engine with a peptide tolerance of 5 ppm and fragment ion tolerance of 30 mmu. Tryptic peptides were accepted with one missed cleavage, oxidation on methionine was set as a variable modification, and methylthiolation on cysteine was set as a fixed modification. Fixed Value PSM Validator was used for PSM validation.

## Data availability

The sub-tomogram averages can be accessed with the following accession codes EMD-17391, EMD-17468 and EMD-17467. Representative tomograms and the raw tilt series have been uploaded to the Electron Microscopy Public Image Archive (EMPIAR) and can be accessed with the deposition EMPIAR-11557. Mass-spectrometry data has been uploaded to PRIDE under project accession PXD042571.

## Acknowledgments

We thank all members of the Barandun laboratory and the Carlson laboratory for the helpful discussions. Further, we thank Michael Hall and Camilla Holmlund for their help with cryo-EM data collection. The electron microscopy data was collected at the Umeå Core Facility for Electron Microscopy, a node of the Cryo-EM Swedish National Facility, funded by the Knut and Alice Wallenberg, Family Erling Persson and Kempe Foundations, SciLifeLab, Stockholm University, and Umeå University. The authors also thank the Proteomics Core Facility at Sahlgrenska Academy, University of Gothenburg, for the proteomic analysis. H.S. is supported by the MSCA fellowship “MsInfection” (Grant agreement ID: 101033469). N.J. is supported by an Integrated Structural Biology fellowship from the Kempe Foundation (JCK-1918). J.B. acknowledges funding from the Swedish Research Council (2019-02011), the European Research Council (ERC Starting Grant PolTube 948655), the SciLifeLab National Fellows program, and MIMS.

## Author contributions

H.S. and J.B. conceived the study. N.J. performed germination experiments and analysis and, together with K.E., helped H.S with electron microscopy work. H.S., with the help of L.A.C., performed the Cryo-ET analysis. K.E. performed protein purification work. All authors interpreted the results, wrote, and edited the manuscript.

## References

Bacela-Spychalska K, Wróblewski P, Mamos T, Grabowski M, Rigaud T, Wattier R, Rewicz T, Konopacka A & Ovcharenko M (2018) Europe-wide reassessment of Dictyocoela (Microsporidia) infecting native and invasive amphipods (Crustacea): molecular versus ultrastructural traits. Sci Rep 8: 8945

Barandun J, Hunziker M, Vossbrinck CR & Klinge S (2019) Evolutionary compaction and adaptation visualized by the structure of the dormant microsporidian ribosome. Nat Microbiol 4: 1798–1804

Bouzahzah B, Nagajyothi F, Ghosh K, Takvorian PM, Cali A, Tanowitz HB & Weiss LM (2010) Interactions of encephalitozoon cuniculi polar tube proteins. Infect Immun 78: 2745–2753

Vande Broek A, Lambrecht M, Eggermont K & Vanderleyden J (1999) Auxins upregulate expression of the indole-3-pyruvate decarboxylase gene in Azospirillum brasilense. J Bacteriol 181: 1338–42

Byers B (1971) Chick embryo ribosome crystals: analysis of bonding and functional activity in vitro. Proc Natl Acad Sci U S A 68: 440–444

Campbell SE, Williams TA, Yousuf A, Soanes DM, Paszkiewicz KH & Williams BAP (2013) The genome of Spraguea lophii and the basis of host-microsporidian interactions. PLoS Genet 9: e1003676

Casanova M, Lopez-Ribot JL, Monteagudo C, Llombart-Bosch A, Sentandreu R & Martinez JP (1992) Identification of a 58-kilodalton cell surface fibrinogen-binding mannoprotein from Candida albicans. Infect Immun 60: 4221–4229

Castaño-Díez D, Kudryashev M, Arheit M & Stahlberg H (2012) Dynamo: a flexible, user-friendly development tool for subtomogram averaging of cryo-EM data in high-performance computing environments. J Struct Biol 178: 139–151

Chang R, Davydov A, Jaroenlak P, Budaitis B, Ekiert DC, Bhabha G & Prakash M (2023) Energetics of the Microsporidian Polar Tube Invasion Machinery. bioRxiv Prepr Serv Biol: 2023.01.17.524456

Chen M, Dai W, Sun SY, Jonasch D, He CY, Schmid MF, Chiu W & Ludtke SJ (2017) Convolutional neural networks for automated annotation of cellular cryo-electron tomograms. Nat Methods 14: 983–985

Corradi N (2015) Microsporidia: Eukaryotic Intracellular Parasites Shaped by Gene Loss and Horizontal Gene Transfers. Annu Rev Microbiol 69: 167–183

Cummings RD & Etzler ME (2009) R-type Lectins Cold Spring Harbor Laboratory Press

Dean P, Sendra KM, Williams TA, Watson AK, Major P, Nakjang S, Kozhevnikova E, Goldberg A V, Kunji ERS, Hirt RP, et al (2018) Transporter gene acquisition and innovation in the evolution of Microsporidia intracellular parasites. Nat Commun 9: 1709

Ehrenbolger K, Jespersen N, Sharma H, Sokolova YY, Tokarev YS, Vossbrinck CR & Barandun J (2020) Differences in structure and hibernation mechanism highlight diversification of the microsporidian ribosome. PLoS Biol 18: 1–15

Fakhree MAA, Blum C & Claessens MMAE (2019) Shaping membranes with disordered proteins. Arch Biochem Biophys 677: 108163

Fayet M, Prybylski N, Collin M-L, Peyretaillade E, Wawrzyniak I, Belkorchia A, Akossi RF, Diogon M, Alaoui H El, Polonais V, et al (2023) Identification and localization of polar tube proteins in the extruded polar tube of the microsporidian Anncaliia algerae. doi:10.21203/rs.3.rs-2507613/v1 [PREPRINT]

Franken LE, Oostergetel GT, Pijning T, Puri P, Arkhipova V, Boekema EJ, Poolman B & Guskov A (2017) A general mechanism of ribosome dimerization revealed by single-particle cryo-electron microscopy. Nat Commun 8: 722

Franzen C (2004) Microsporidia: how can they invade other cells? Trends Parasitol 20: 275–279

Fries I, Martín R, Meana A, García-Palencia P & Higes M (2006) Natural infections of Nosema ceranae in European honey bees. J Apic Res 45: 230–233

Frixione E, Ruiz L, Santillán M, de Vargas L V, Tejero JM & Undeen AH (1992) Dynamics of polar filament discharge and sporoplasm expulsion by microsporidian spores. Cell Motil Cytoskelet 22: 38–50

Hagen WJH, Wan W & Briggs JAG (2017) Implementation of a cryoelectron tomography tilt-scheme optimized for high resolution subtomogram averaging. J Struct Biol 197: 191–198

Han B, Ma Y, Tu V, Tomita T, Mayoral J, Williams T, Horta A, Huang H & Weiss LM (2019) Microsporidia Interact with Host Cell Mitochondria via Voltage-Dependent Anion Channels Using Sporoplasm Surface Protein 1. MBio 10

Han B, Polonais V, Sugi T, Yakubu R, Takvorian PM, Cali A, Maier K, Long M, Levy M, Tanowitz HB, et al (2017) The role of microsporidian polar tube protein 4 (PTP4) in host cell infection. PLoS Pathog 13: 1–28

Han B, Takvorian PM & Weiss LM (2020) Invasion of Host Cells by Microsporidia. Front Microbiol 11: 172

Han B & Weiss LM (2017) Microsporidia: Obligate Intracellular Pathogens Within the Fungal Kingdom. Microbiol Spectr 5

Heinz E, Williams TA, Nakjang S, Noël CJ, Swan DC, Goldberg A V, Harris SR, Weinmaier T, Markert S, Becher D, et al (2012) The genome of the obligate intracellular parasite Trachipleistophora hominis: new insights into microsporidian genome dynamics and reductive evolution. PLoS Pathog 8: e1002979

Hounsell EF, Davies MJ & Renouf D V. (1996) O-linked protein glycosylation structure and function. Glycoconj J 13: 19–26

Ishida T & Kinoshita K (2007) PrDOS: prediction of disordered protein regions from amino acid sequence. Nucleic Acids Res 35: W460–W464

Jaroenlak P, Cammer M, Davydov A, Sall J, Usmani M, Liang F-X, Ekiert DC & Bhabha G (2020) 3-Dimensional organization and dynamics of the microsporidian polar tube invasion machinery. PLOS Pathog 16: e1008738

Jespersen N, Ehrenbolger K, Winiger RR, Svedberg D, Vossbrinck CR & Barandun J (2022a) Structure of the reduced microsporidian proteasome bound by PI31-like peptides in dormant spores. Nat Commun 13: 6962

Jespersen N, Monrroy L & Barandun J (2022b) Impact of Genome Reduction in Microsporidia. Exp Suppl 114: 1–42

Karpova EA & Gillet R (2018) The Structural and Functional Organization of Ribosomal Compartment in the Cell: A Mystery or a Reality? Trends Biochem Sci 43: 938–950

Keohane EM, Orr GA, Zhang HS, Takvorian PM, Cali A, Tanowitz HB, Wittner M & Weiss LM (1998) The molecular characterization of the major polar tube protein gene from Encephalitozoon hellem, a microsporidian parasite of humans. Mol Biochem Parasitol 94: 227–236

Kremer JR, Mastronarde DN & McIntosh JR (1996) Computer visualization of three-dimensional image data using IMOD. J Struct Biol 116: 71–76

Kurtti TJ, Munderloh UG & Noda H (1990) Vairimorpha necatrix: Infectivity for and development in a lepidopteran cell line. J Invertebr Pathol 55: 61–68

Kusamrarn T, Sobhon P & Bailey GB (1975) The mechanism of formation of inhibitor-induced ribosome helices in Entamoeba invadens. J Cell Biol 65: 529–539

Lin B, Qing X, Liao J & Zhuo K (2020) Role of Protein Glycosylation in Host-Pathogen Interaction. Cells 9: 1022

Liu H, Li M, Cai S, He X, Shao Y & Lu X (2016) Ricin-B-lectin enhances microsporidia Nosema bombycis infection in BmN cells from silkworm Bombyx mori. Acta Biochim Biophys Sin 48: 1050–1057

Liu Y-T, Zhang H, Wang H, Tao C-L, Bi G-Q & Zhou ZH (2022) Isotropic reconstruction for electron tomography with deep learning. Nat Commun 13: 6482

Lv Q, Wang L, Fan Y, Meng X, Liu K, Zhou B, Chen J, Pan G, Long M & Zhou Z (2020) Identification and characterization a novel polar tube protein (NbPTP6) from the microsporidian Nosema bombycis. Parasit Vectors 13: 475

Maheshwari AJ, Sunol AM, Gonzalez E, Endy D & Zia RN (2023) Colloidal Physics Modeling Reveals How Per-Ribosome Productivity Increases with Growth Rate in Escherichia coli. MBio 14

Major P, Sendra KM, Dean P, Williams TA, Watson AK, Thwaites DT, Embley TM & Hirt RP (2019) A new family of cell surface located purine transporters in Microsporidia and related fungal endoparasites. Elife 8: e47037

Matzov D, Bashan A, Yap M-NF & Yonath A (2019) Stress response as implemented by hibernating ribosomes: a structural overview. FEBS J 286: 3558–3565

McLaren M, Gil-Diez P, Isupov MN, Conners R, Gambelli L, Gold V, Walter A, Connell SR, Williams B & Daum B (2022) In situ structure of a dimeric hibernating ribosome from a eukaryotic intracellular pathogen. bioRxiv: 2022.04.29.490036 [PREPRINT]

Milligan RA & Unwin PN (1982) In vitro crystallization of ribosomes from chick embryos. J Cell Biol 95: 648–653

Mitchell MJ & Cali A (1994) Vairimorpha necatrix (Microsporida: Burenellidae) Affects Growth and Development of Heliothis zea (Lepidoptera: Noctuidae) Raised at Various Temperatures. J Econ Entomol 87: 933–940

Morimoto T, Blobel G & Sabatini DD (1972a) Ribosome crystallization in chicken embryos. I. Isolation, characterization, and in vitro activity of ribosome tetramers. J Cell Biol 52: 338–354

Morimoto T, Blobel G & Sabatini DD (1972b) Ribosome crystallization in chicken embryos. II. Conditions for the formation of ribosome tetramers in vitro. J Cell Biol 52: 355–366

Nakjang S, Williams TA, Heinz E, Watson AK, Foster PG, Sendra KM, Heaps SE, Hirt RP & Martin Embley T (2013) Reduction and expansion in microsporidian genome evolution: new insights from comparative genomics. Genome Biol Evol 5: 2285–2303

Nicholson D, Salamina M, Panek J, Helena-Bueno K, Brown CR, Hirt RP, Ranson NA & Melnikov S V (2022) Adaptation to genome decay in the structure of the smallest eukaryotic ribosome. Nat Commun 13: 591

Nogueira PM, Assis RR, Torrecilhas AC, Saraiva EM, Pessoa NL, Campos MA, Marialva EF, Ríos-Velasquez CM, Pessoa FA, Secundino NF, et al (2016) Lipophosphoglycans from Leishmania amazonensis Strains Display Immunomodulatory Properties via TLR4 and Do Not Affect Sand Fly Infection. PLoS Negl Trop Dis 10: e0004848

Palenzuela O, Redondo MJ, Cali A, Takvorian PM, Alonso-Naveiro M, Alvarez-Pellitero P & Sitjà-Bobadilla A (2014) A new intranuclear microsporidium, Enterospora nucleophila n. sp., causing an emaciative syndrome in a piscine host (Sparus aurata), prompts the redescription of the family Enterocytozoonidae. Int J Parasitol 44: 189–203

Petri Jr WA, Haque R & Mann BJ (2002) The bittersweet interface of parasite and host: lectin-carbohydrate interactions during human invasion by the parasite Entamoeba histolytica. Annu Rev Microbiol 56: 39–64

Peuvel I, Peyret P, Méténier G, Vivarès CP & Delbac F (2002) The microsporidian polar tube: Evidence for a third polar tube protein (PTP3) in Encephalitozoon cuniculi. Mol Biochem Parasitol 122: 69–80

Prates ET, Guan X, Li Y, Wang X, Chaffey PK, Skaf MS, Crowley MF, Tan Z & Beckham GT (2018) The impact of O-glycan chemistry on the stability of intrinsically disordered proteins. Chem Sci 9: 3710–3715

Prybylski N, Fayet M, Dubuffet A, Delbac F, Kocer A, Gardarin C, Michaud P, El Alaoui H & Dubessay P (2022) Ricin B lectin-like proteins of the microsporidian Encephalitozoon cuniculi and Anncaliia algerae are involved in host-cell invasion. Parasitol Int 87: 102518

Rauscher S & Pomès R (2017) The liquid structure of elastin. Elife 6: e26526

Sardinha-Silva A, Mendonça-Natividade FC, Pinzan CF, Lopes CD, Costa DL, Jacot D, Fernandes FF, Zorzetto-Fernandes AL V, Gay NJ, Sher A, et al (2019) The lectin-specific activity of Toxoplasma gondii microneme proteins 1 and 4 binds Toll-like receptor 2 and 4 N-glycans to regulate innate immune priming. PLoS Pathog 15: e1007871

Schindelin J, Arganda-Carreras I, Frise E, Kaynig V, Longair M, Pietzsch T, Preibisch S, Rueden C, Saalfeld S, Schmid B, et al (2012) Fiji: an open-source platform for biological-image analysis. Nat Methods 9: 676–682

Takvorian PM, Han B, Cali A, Rice WJ, Gunther L, Macaluso F & Weiss LM (2020) An Ultrastructural Study of the Extruded Polar Tube of Anncaliia algerae (Microsporidia). J Eukaryot Microbiol 67: 28–44

Tristram-Nagle S & Nagle JF (2004) Lipid bilayers: thermodynamics, structure, fluctuations, and interactions. Chem Phys Lipids 127: 3–14

Troemel ER & Becnel JJ (2015) Genome analysis and polar tube firing dynamics of mosquito-infecting microsporidia. Fungal Genet Biol 83: 41–44

Troemel ER, Félix M-A, Whiteman NK, Barrière A & Ausubel FM (2008) Microsporidia Are Natural Intracellular Parasites of the Nematode Caenorhabditis elegans. PLoS Biol 6: e309

Wadi L & Reinke AW (2020) Evolution of microsporidia: An extremely successful group of eukaryotic intracellular parasites. PLoS Pathog 16: e1008276

Weidner E (1976) The microsporidian spore invasion tube: The ultrastructure, isolation, and characterization of the protein comprising the tube. J Cell Biol 71: 23–34

Wiśniewski JR, Zougman A, Nagaraj N & Mann M (2009) Universal sample preparation method for proteome analysis. Nat Methods 6: 359–362

Xu Y, Takvorian PM, Cali A, Orr G & Weiss LM (2004) Glycosylation of the major polar tube protein of Encephalitozoon hellem, a microsporidian parasite that infects humans. Infect Immun 72: 6341–6350

Xu Y & Weiss LM (2005) The microsporidian polar tube: A highly specialised invasion organelle. Int J Parasitol 35: 941–953

Yachnis AT, Berg J, Martinez-Salazar A, Bender BS, Diaz L, Rojiani AM, Eskin TA & Orenstein JM (1996) Disseminated microsporidiosis especially infecting the brain, heart, and kidneys. Report of a newly recognized pansporoblastic species in two symptomatic AIDS patients. Am J Clin Pathol 106: 535–543

Yuan F, Alimohamadi H, Bakka B, Trementozzi AN, Day KJ, Fawzi NL, Rangamani P & Stachowiak JC (2021) Membrane bending by protein phase separation. Proc Natl Acad Sci U S A 118

Zheng SQ, Palovcak E, Armache J-P, Verba KA, Cheng Y & Agard DA (2017) MotionCor2: anisotropic correction of beam-induced motion for improved cryo-electron microscopy. Nat Methods 14: 331–332

