## Supplementary Data file for "Ribosome clustering and surface layer reorganization in the microsporidian host-invasion apparatus"

**Supplementary Table 1.** Cryo-ET data collection parameters.

**Supplementary Table 2.** Mass spectrometry results of a PTP3 affinity-purified sample.

**Supplementary Figure 1.** Tracking polar tube eversion to understand germination dynamics and tube length.

**Supplementary Figure 2**. Representative tomograms depicting empty or sporoplasm-packed germinated polar tubes.

**Supplementary Figure 3**. Subtomogram averaging workflow.

**Supplementary Figure 4**. Identification of PTP6 and its isoforms in PTP3 pulldowns.

**Supplementary_video_v1.** Tomographic volume of a ribosome-filled-germinated polar tube overlaid with the corresponding segmentation.

**Supplementary Table 1. Cryo-ET data collection parameters.**

| **Data collection parameters** | |
| --- | --- |
| Microscope | Titan Krios G2 |
| Voltage (kV) | 300 |
| Camera | Gatan K2 |
| Energy filter | Yes, BioQuantum |
| Magnification | 64000x |
| Total Electron dosage (e–/Å^2^ ) | 110 to 120 |
| Defocus range (μm) | -1.5 to 5 |
| Tilt range (°) | -60 to +60 |
| Tilt angle increment (°) | 2/3 |
| Object pixel size (Å) | 2.173 |
| Number of tomograms | 50 |

**Supplementary Table 2. Mass spectrometry results of a PTP3 affinity-purified sample.** The list of mass spectrometry hits is arranged according to peptide-spectrum match (PSM). The number of consecutive histidines (3xHis, 4xHis, 7xHis) in the protein is listed. PTP3 is the only protein with seven consecutive histidines.

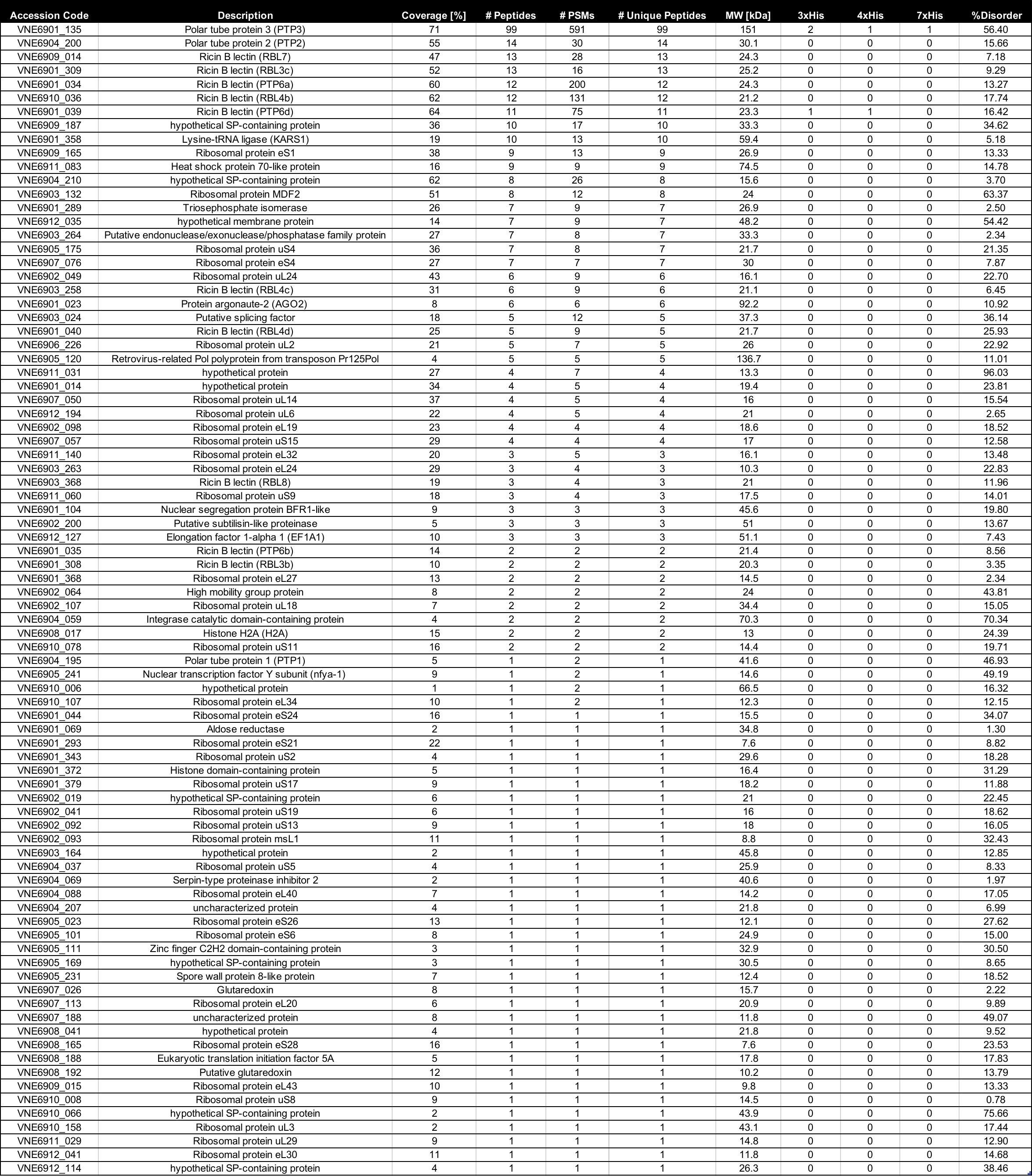

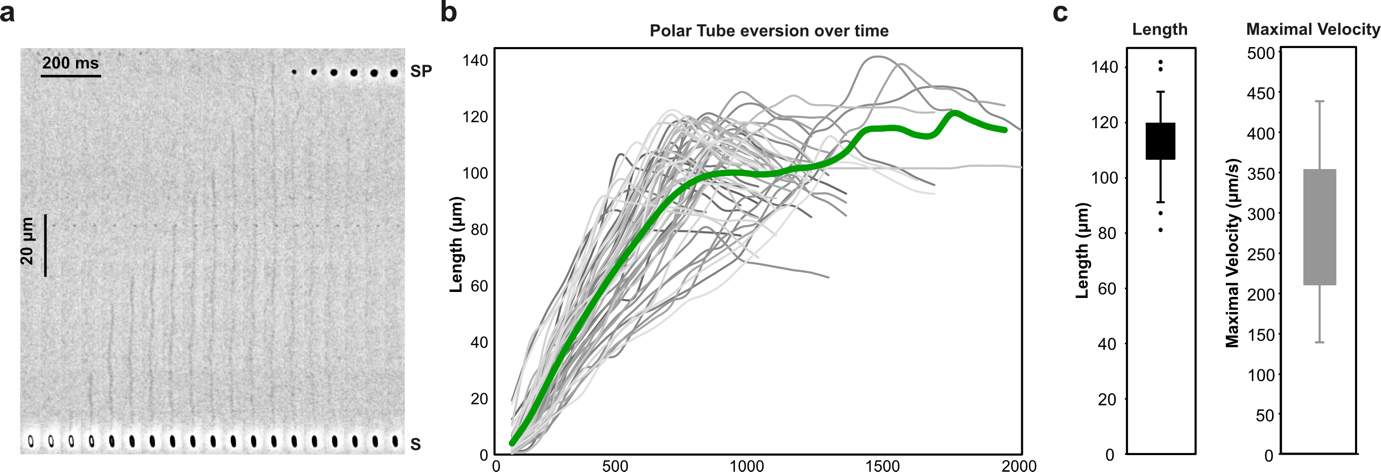

**Supplementary Figure 1. Tracking polar tube eversion to understand germination dynamics and tube length.** (**a**) A kymograph obtained via live light microscopy analysis of polar tube firing events from *Vairimorpha necatrix.* (**b**) Length over time diagrams of all analyzed polar tube eversion events. The average length over time is colored in green. (**c**) Bar plots of polar tube length and maximum velocity distribution.

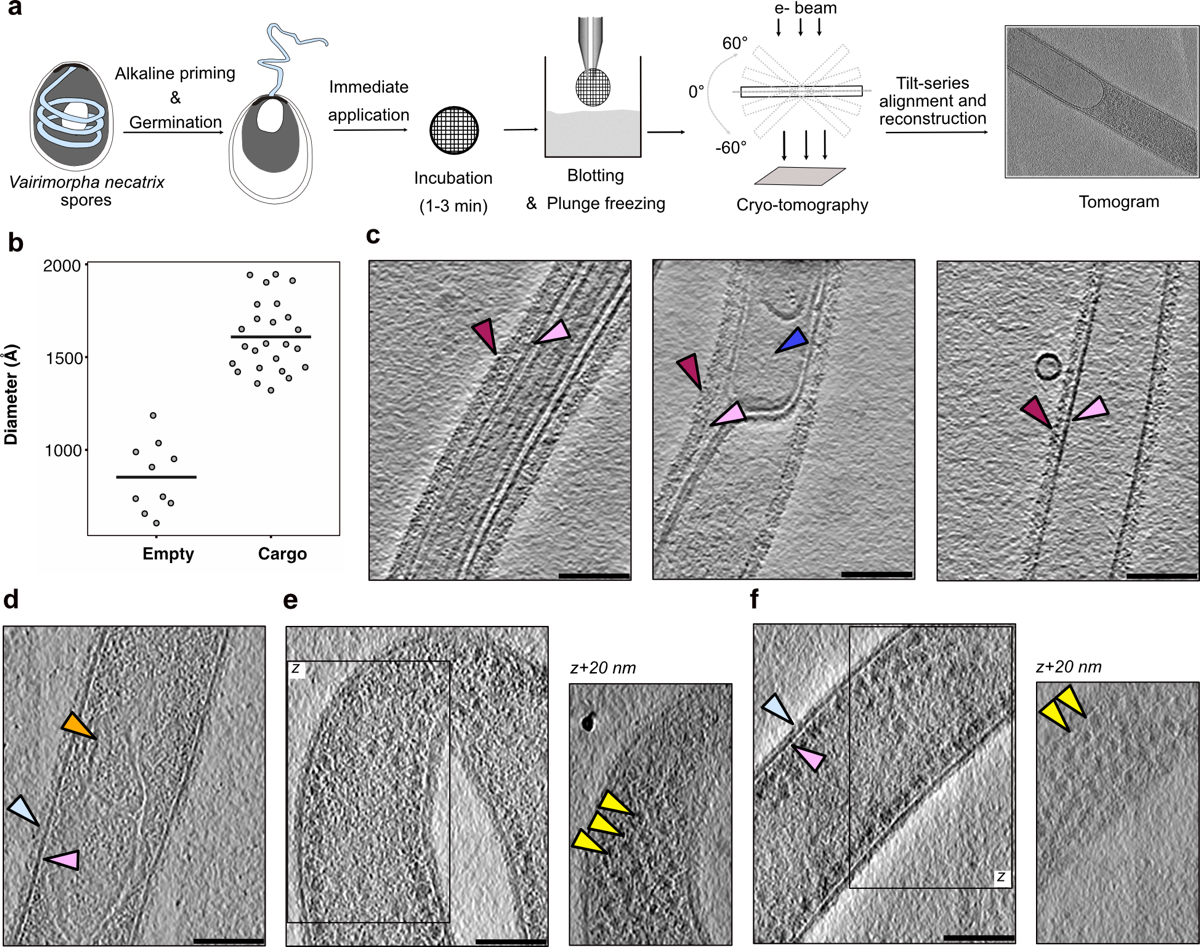

**Supplementary Figure 2**. **Representative tomograms depicting empty or sporoplasm-packed germinated polar tubes.** (**a**) A schematic representation of the methodology for on-grid freezing and collecting tomograms of germinated polar tubes. (**b**) A graph representing the internal diameter of polar tubes (*PTempty* & *PTcargo*) visualized using cryo-ET. Each dot represents one tube, and the line represents the mean diameter. **(c**) Representative tomograms of *PTempty*, or polar tubes devoid of cellular cargo or filled with electron-lucent material. The central section of a tomogram is shown with regions of interest indicated with arrows (magenta for the outer wall, pink for the lipid bilayer, and blue for vesicles). (**d-f**) Representative tomograms from *PTcargo*, or polar tubes filled with cellular cargo where (e-f) contained ribosome spirals inside tubes. The central section of a tomogram is presented, and regions of interest are indicated with arrows (light blue for the outer tube wall, pink for the lipid bilayer, and yellow for ribosomes). For (e-f), additional views corresponding to the boxed regions are also presented.

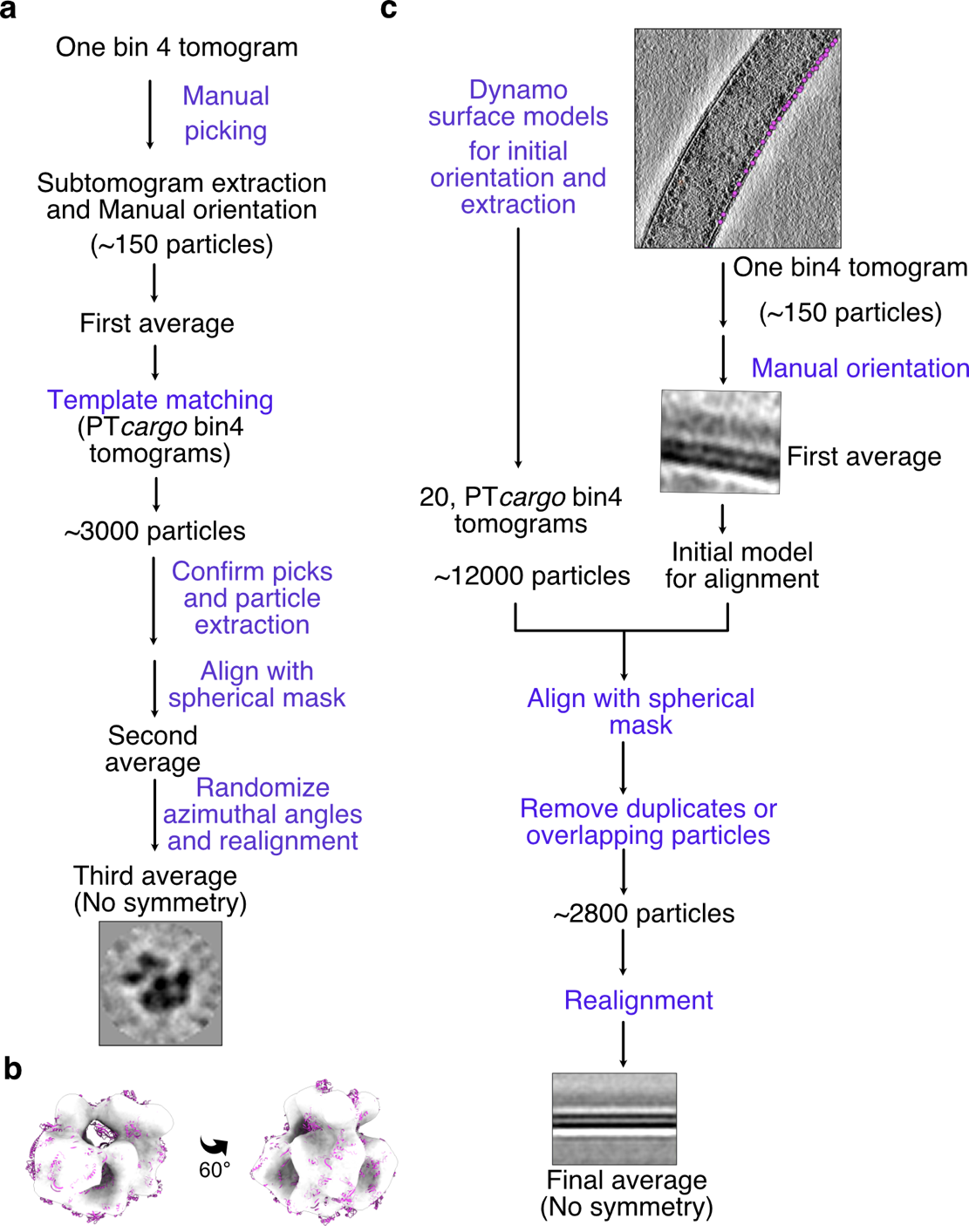

**Supplementary Figure 3**. **Subtomogram averaging workflow. (a)** Schematic workflow of the subtomogram averaging procedure to generate the ribosome volume. **(b)** Two 60°-related views of ribosome reconstruction (transparent white), fitted with the structure of the *V. necatrix* ribosome (PDB ID: 6RM3, magenta). **(c)** Schematic workflow of the subtomogram averaging procedure used to create the reconstructions of the segments of polar tube outer layers. The scheme is shown for cargo-filled tubes, and a similar methodology was used for empty tubes.

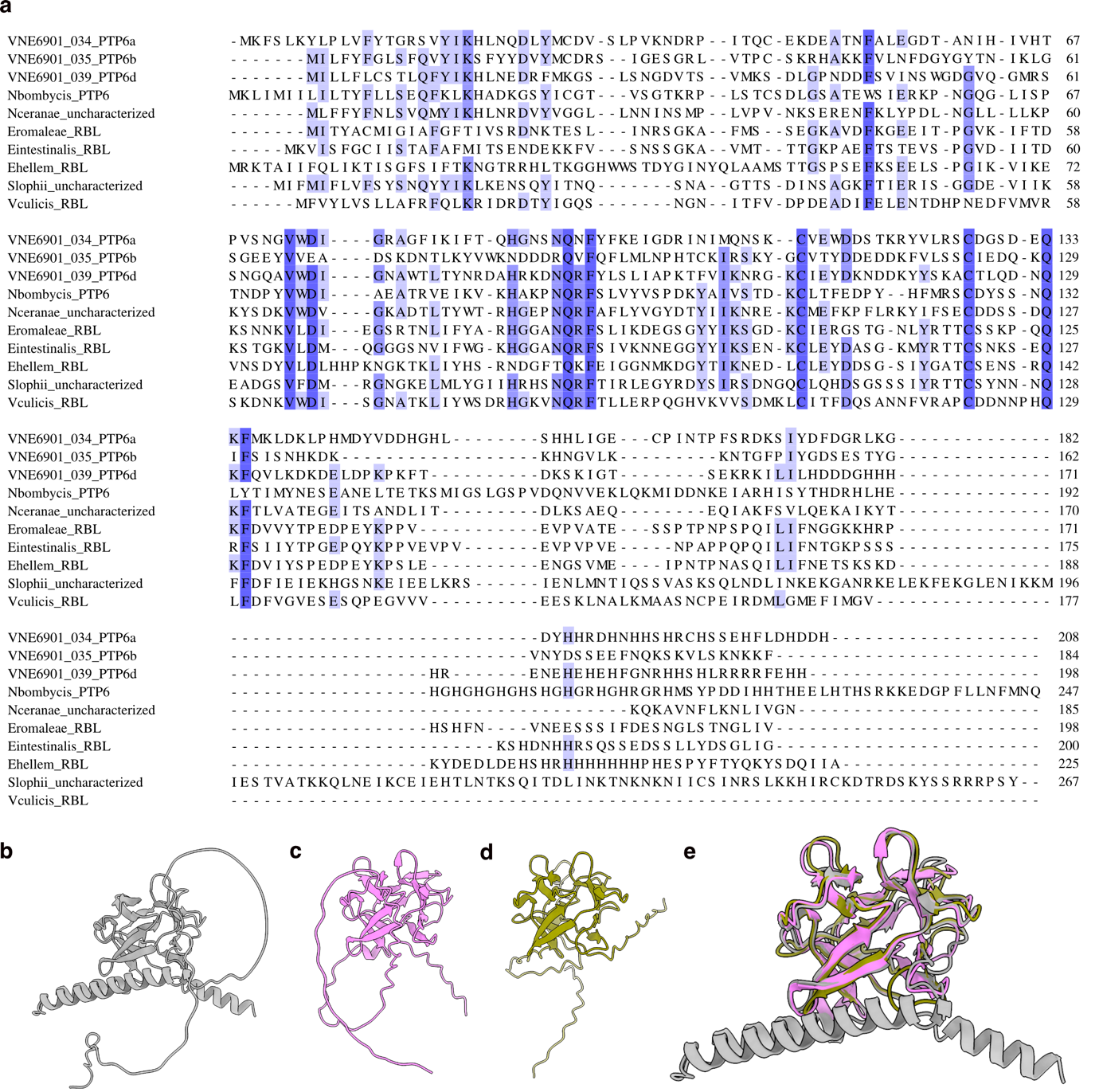

**Supplementary Figure 4**. **Identification of PTP6 and its isoforms in PTP3 pulldowns. (a)** Alignment of VNE6901_039 and VNE6901_034 with PTP6 homologs from *Nosema bombycis* (R0MBR8_NOSB1), *Vavraia culicis* (L2GVW3_VAVCU), *Nosema ceranae* (C4V7Y1_NOSC), *Spraguea lophii* (S7XK85_SPRLO), *Encephalitozoon hellem* (I6TKU6_ENCHA), *Encephalitozoon romaleae* (I7AT09_ENCRO), *Encephalitozoon intestinalis* (E0S8R2_ENCIT). Protein sequences were retrieved from Uniprot and aligned using Muscle followed by visualization using Jalview. **(b-d)** Alphfold models for *N. bombycis* PTP6 (b), VNE6901_039 (c) and VNE6901_034 (d). **(e)** Overlay of predicted PTP6 models from (b-d) shown at 90° rotation. Regions predicted with low confidence have been excluded for clarity.
